# ZonationR: An R interface to the Zonation software for reproducible spatial conservation prioritisation workflows

**DOI:** 10.64898/2026.04.28.720523

**Authors:** Thiago Cavalcante, Bruno Ribeiro, Karlo Guidoni-Martins, Heini Kujala

## Abstract

Systematic conservation planning provides a science-based framework for defining conservation goals and supporting transparent spatial decision-making under limited resources. Within this framework, spatial conservation prioritisation tools are widely used to identify areas of high biodiversity value by integrating information on species distributions, connectivity, costs, and other factors into spatially explicit recommendations. Zonation is one of the leading software tools in this field, producing hierarchical priority rankings of landscapes based on conservation value. However, its standard workflow typically relies on manual steps for data preparation, execution, and post-processing, which can become inefficient and difficult to reproduce when multiple scenarios are analysed, limiting accessibility and broader uptake. We introduce *ZonationR*, an R package that provides a streamlined interface to the Zonation software, enabling fully reproducible and automated spatial prioritisation workflows. The package integrates the entire analysis pipeline, encompassing input preparation, execution of Zonation, and post-processing, while supporting both single-variant and multi-variant workflows. *ZonationR* also provides tools to explore and interpret outputs, including priority maps, feature performance curves, cost summaries, feature representation metrics, and similarity assessments between prioritisation solutions. By linking directly to the original Zonation engine, the package enables users to benefit from ongoing methodological developments in Zonation and access its functionality through a transparent, script-based workflow, thereby reducing technical barriers to running and understanding spatial prioritisation analyses. Beyond these advantages, its integration within the R environment supports iterative testing of conservation scenarios and more rigorous assessment of methodological decisions, while facilitating seamless connections with wider ecological workflows (e.g., species distribution modelling). As conservation planning increasingly relies on large, complex, multi-source datasets and integrative approaches, such integration is essential for enabling robust, transparent, and reproducible decision-making across spatial scales.

## 1. Background

Systematic conservation planning is a structured, science-based approach for setting clear conservation goals and guiding transparent decision-making (Margules and Pressey, 2000). Within this framework, spatial conservation prioritisation tools help identify optimal areas to protect biodiversity under limited resources (Sarkar et al., 2006; Giakoumi et al., 2025). These tools differ in their algorithms, approaches, and the conservation objectives they are designed to address (Moilanen et al., 2009; Giakoumi et al., 2025). They integrate information on species distributions, habitat connectivity, and costs (e.g. land acquisition expenses or opportunity costs) to translate systematic conservation planning principles into practical, spatially explicit guidance. In doing so, they provide a rigorous, quantitative foundation for identifying priority areas where conservation efforts can have the greatest impact.

Zonation is one of the leading conservation planning software tools supporting this process, offering robust algorithms and flexible workflows for generating spatially explicit priority maps (Moilanen et al., 2005; Moilanen et al., 2022). It ranks locations across the landscape based on spatial data on biodiversity value, connectivity, costs, among other factors, providing spatial outputs and decision-support information for both policy development and scientific research (Arponen et al., 2005; Lehtomäki and Moilanen, 2013). While Zonation is a powerful and widely used tool, its standard workflow typically requires manual preparation of input files, execution through a graphical interface or command line, and post-processing carried out separately. This workflow becomes increasingly inefficient when multiple scenarios or repeated runs are required (which is often the case), as these manual steps limit transparency, consistency, and reproducibility. As a result, Zonation’s workflow complexity can limit its uptake among ecologists as well as the broader community of applied researchers and educators.

*ZonationR* addresses these limitations by providing a seamless R interface to Zonation. The package automates the creation of input files, calls Zonation to perform the spatial prioritisation analysis, and imports outputs for streamlined post-processing, visualization, and output comparison. By integrating these steps into a single workflow, *ZonationR* enhances reproducibility, facilitates automation, and simplifies the sharing of analysis pipelines, making the entire process more efficient and broadly accessible. Importantly, integrating Zonation with R enables its direct incorporation into broader analytical pipelines, including data gathering and processing, species distribution modelling, and scenario-based analyses, thereby supporting more iterative and flexible conservation planning workflows. It also enables users to compare different prioritisation algorithms and evaluate trade-offs between alternative conservation objectives, which is essential for understanding how objective formulation affects conservation outcomes (Cavalcante et al., 2025).

The package interfaces directly with the original Zonation software, maintaining consistency with its well-tested and widely validated implementation (C++), while avoiding duplication of complex algorithmic code in a less computationally efficient R environment. This design enables users to benefit from ongoing methodological developments in Zonation and to access its functionality through a transparent, script-based workflow, thereby lowering the technical barrier to running and understanding spatial prioritisation workflows.

## 2. Package features and workflow

Spatial conservation prioritisation in Zonation is based on an optimized priority ranking of landscape cells according to their conservation value for a set of biodiversity features (e.g., species distributions or habitat types). The algorithm orders cells iteratively according to their marginal contribution to conservation value until it finds the ranking where conservation gains for all included biodiversity features are maximized in the top ranked cell. This process results in a priority rank map where each cell has been ranked from highest to lowest importance for conservation. The framework allows the incorporation of feature weights, conservation costs, ecological condition of locations, existing protected areas and administrative units to guide prioritisation according to conservation objectives. The methodology underlying Zonation has been extensively described and validated in previous studies (see Moilanen et al., 2022). For users experienced with Zonation, it is important to note that *ZonationR* requires Zonation 5 v1.0 or newer, as earlier versions are not supported. Although many of the core principles from older versions, like Zonation 4, are still applied, Zonation 5 is a completely new implementation of the main algorithm and introduces a more intuitive structure for analysis setups, which *ZonationR* leverages to streamline workflows (Moilanen et al., 2022).

The *ZonationR* package provides a modular set of functions designed to streamline the typical workflow stages of spatial conservation prioritisation using the Zonation software (Figure 1). The modular structure also supports step-by-step learning, allowing users to understand how individual components of a spatial prioritisation workflow contribute to the final prioritisation outcomes.

**Figure 1.**
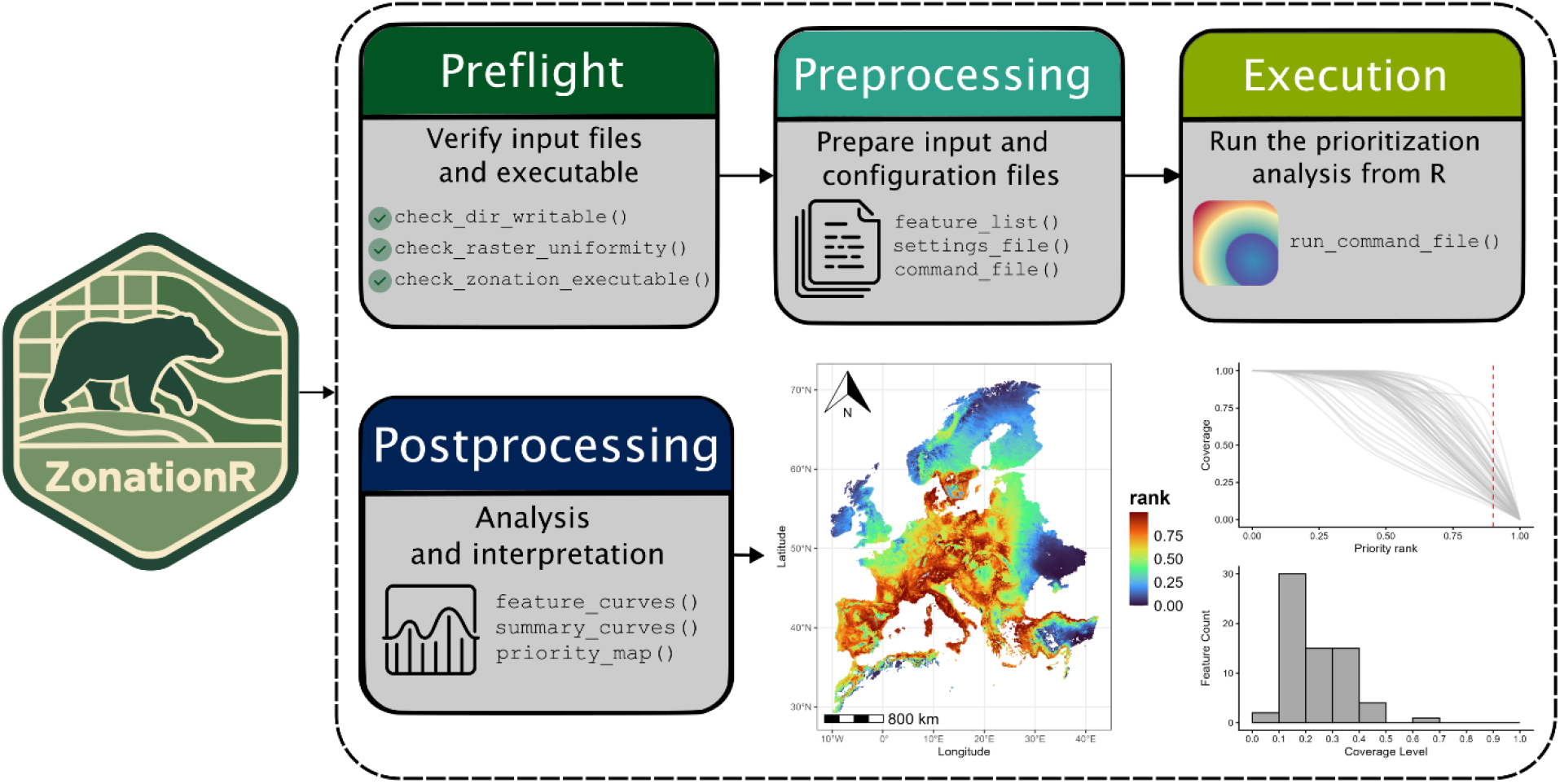
Modular workflow of the *ZonationR* package, illustrating functions that streamline the stages of spatial conservation prioritisation.

The package was developed following best practices for building R packages (Wickham and Bryan, 2023). The *ZonationR* source code is available on GitHub (https://github.com/thiago-cav/ZonationR) and can be installed using the R code below. We also provide a companion data package, available only on GitHub (https://github.com/thiago-cav/Zonation5RData), which contains GeoTIFF layers for workflows with Zonation and *ZonationR*.

~~~
# Install package from CRAN
install.packages(“ZonationR”)
Or install the development version from GitHub:
pak::pak(“thiago-cav/ZonationR”)
# load package
library(ZonationR)
~~~

The *ZonationR* workflow stages include:

- **Preflight**, which involves initial checks such as verifying input files uniformity in terms of resolution, origin, extent and projections, and confirming the availability of the Zonation 5 executable on the system;
- **Preprocessing**, which involves preparing Zonation input files (e.g., feature list) and generating necessary configuration files;
- **Execution**, running the prioritisation analysis from R; and
- **Postprocessing**, where outputs are imported for further analysis and interpretation.

This separation into discrete stages provides a clear and structured framework that facilitates both reproducible analyses and stepwise learning of spatial conservation prioritisation workflows (Table 1).

**Table 1.**
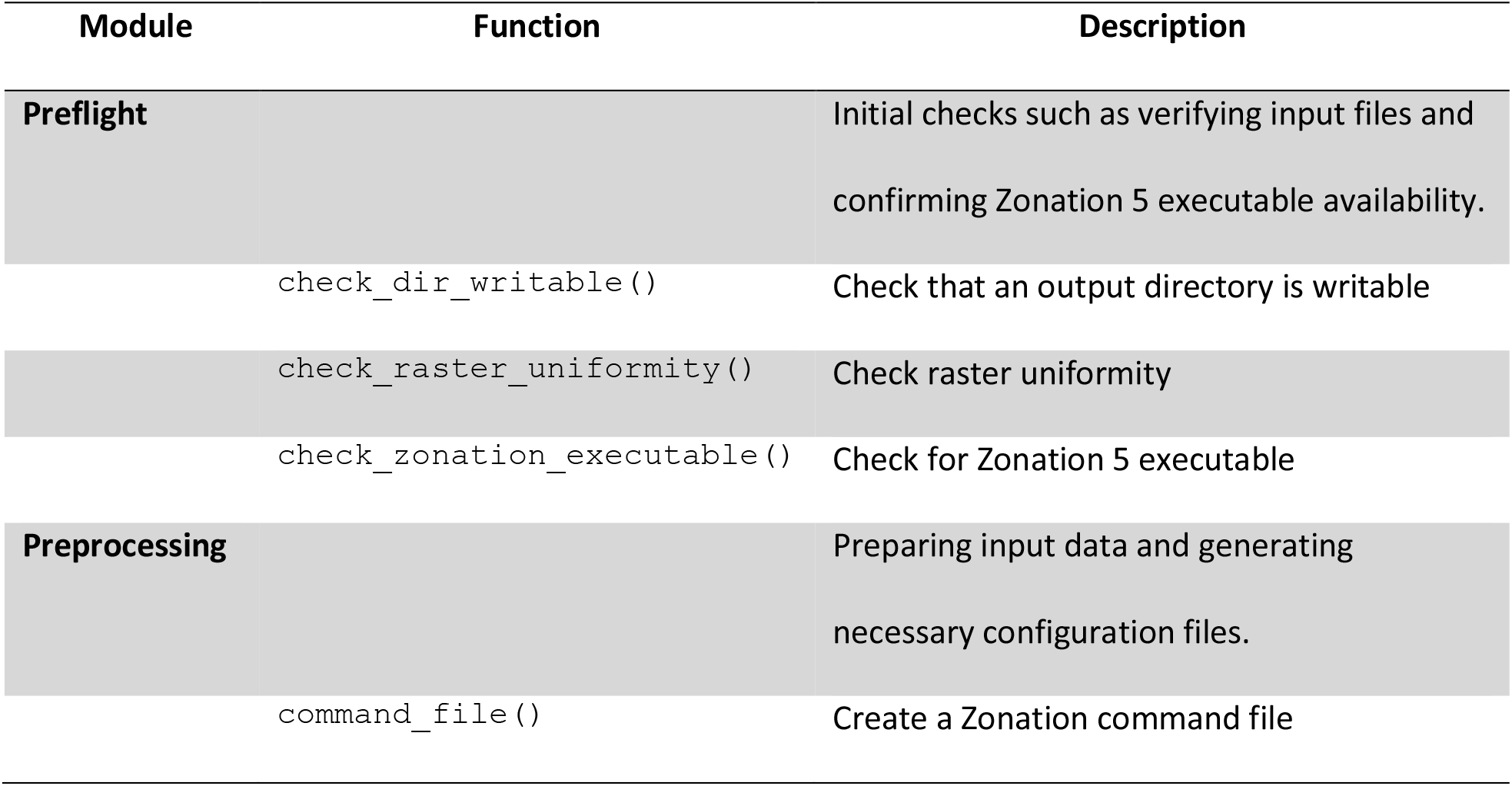
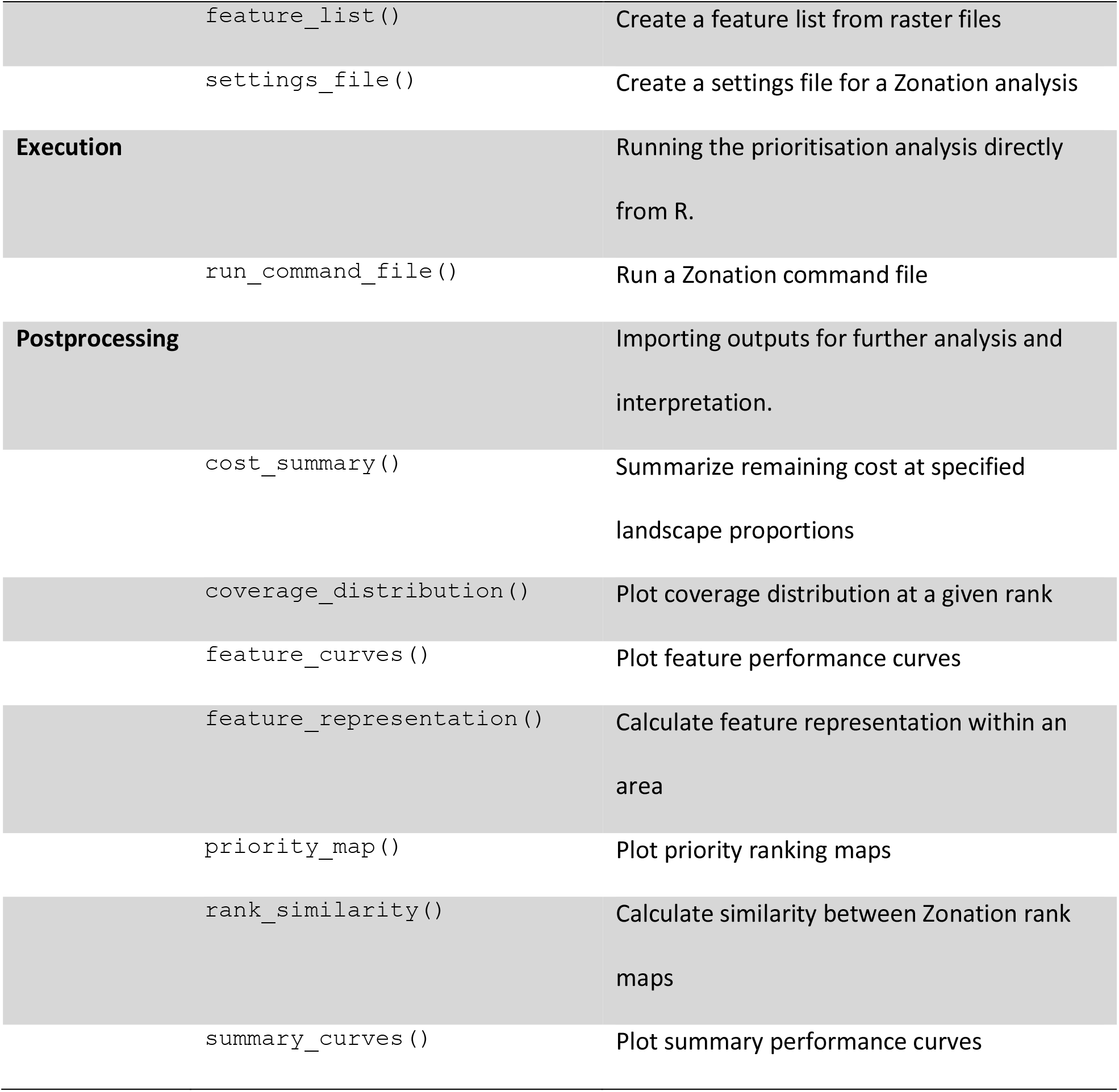
Main stages and functions in the *ZonationR* package.

### 2.1 Configuration and input parameters

The Zonation prioritisation framework is defined through three interacting components: (i) feature specification, (ii) spatial configuration, and (iii) execution control. These components are implemented through the `feature_list()`, `settings_file()`, and `command_file()` functions, which together determine how biodiversity features are represented, how spatial inputs are incorporated, and how prioritisation behaviour is executed.

The feature specification component is defined using the `feature_list()`function, which specifies the biodiversity features included in the analysis. Each entry specifies a feature file, with optional attributes such as weights, output groups, and thresholds that control feature importance, aggregation, and filtering (Table 2). Their application during analysis depends on the activation of corresponding flags in the `command_file()`function (Table 4).

**Table 2.**
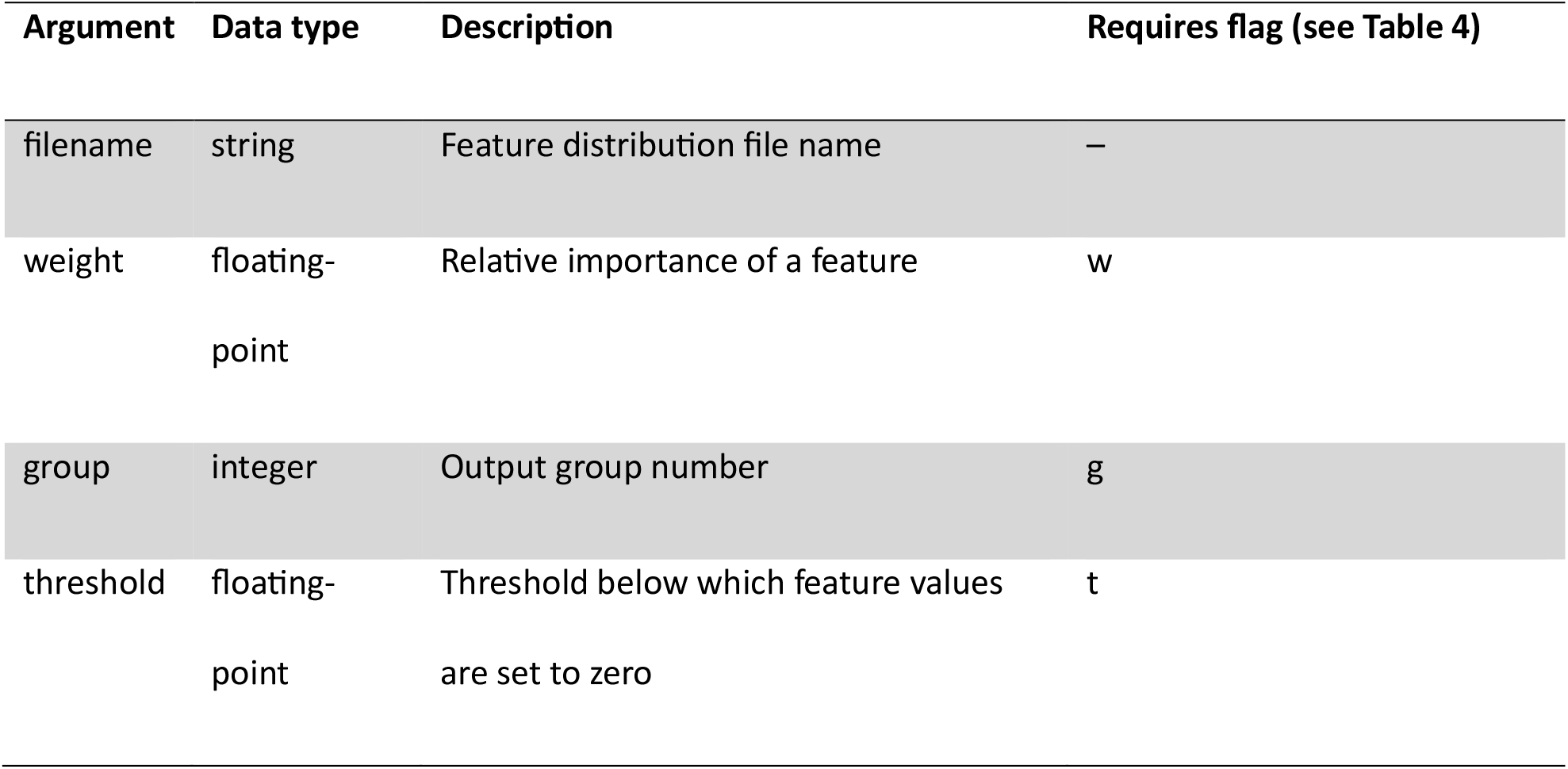
Arguments accepted by the feature_list() function.

| Argument | Data type | Description | Requires flag (see Table 4) |
| --- | --- | --- | --- |
| filename | string | Feature distribution file name | – |
| weight | floating-point | Relative importance of a feature | w |
| group | integer | Output group number | g |
| threshold | floating-point | Threshold below which feature values are set to zero | t |

The spatial configuration component is defined through the `settings_file()` function, which specifies the input data required for the analysis. The feature list file is mandatory, while additional spatial inputs such as analysis masks, hierarchic masks, cost layers, and external solutions are optional (Table 3). These inputs define spatial constraints and cost structures and are only used when activated through the corresponding flags in the `command_file()` function (Table 4).

**Table 3.** Inputs specified in the settings_file() function.

| File parameter | Activation condition (see Table 4) | Required file type |
| --- | --- | --- |
| feature list file | always required | plain text file |
| external solution | --mode=LOAD | floating-point raster |
| analysis area mask | flag a | integer raster |
| hierarchic mask | flag h | integer raster |
| cost layer | flags x, X | floating-point raster |

**Table 4.**
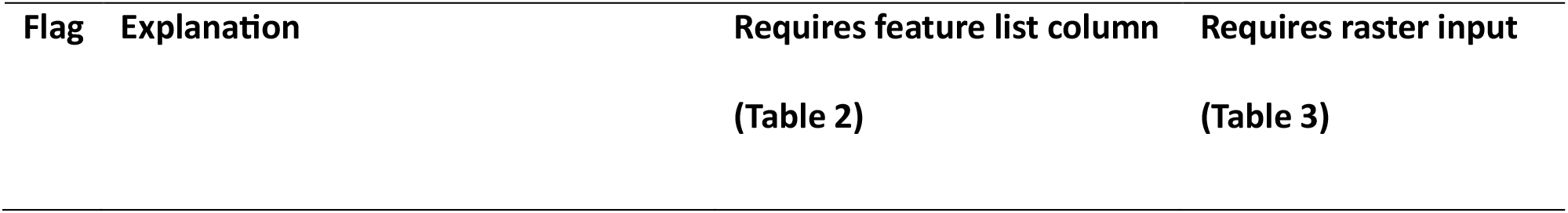
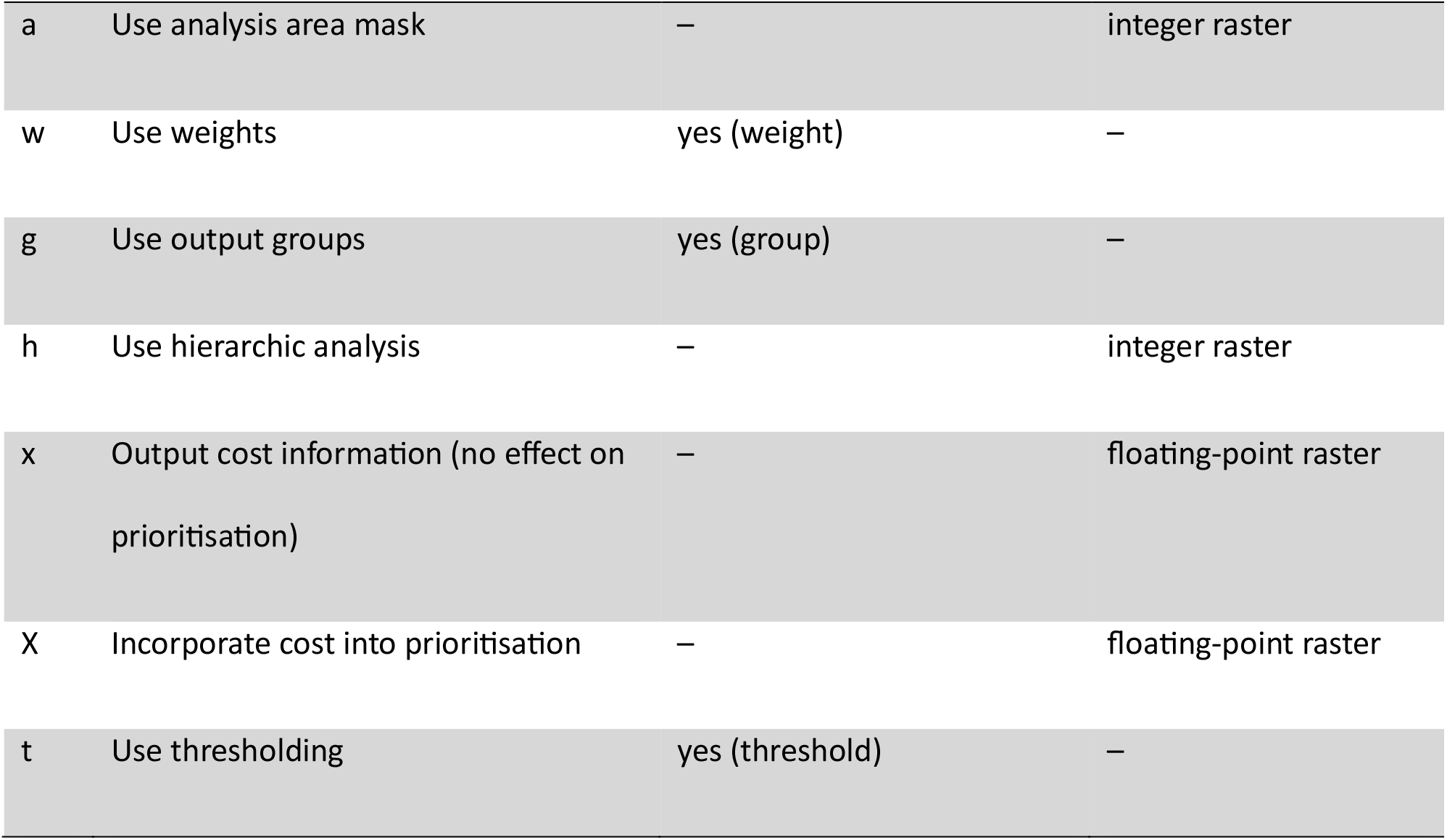
Flags used in the command_file() function.

The execution control component is governed by the `command_file()` function, which defines a set of flags that regulate how feature attributes and spatial inputs are used during prioritisation. Some flags activate feature-level parameters defined in the feature list (Table 2), while others control the use of spatial inputs provided through the settings file (Table 3). Together, these flags determine whether inputs are used for filtering, prioritisation, or output generation (Table 4).

## 3. Application

We demonstrate how *ZonationR* facilitates reproducible spatial conservation prioritisation workflows, including both single-variant analyses and incremental variant workflows. The structured setup below follows the recommended workflow described in the package documentation, allowing users to reproduce examples and adapt them for teaching or training exercises.

### 3.1 Setup and prerequisites

Before running analyses, users should ensure the following:

1. Zonation 5 is installed on the system and accessible from R.
2. Analyses are conducted within a writable project directory (e.g., RStudio projects). This ensures that Zonation can read and write files correctly and that file paths remain consistent.
3. Input biodiversity feature layers are harmonized, i.e., all rasters share the same extent, resolution, and projection.

These requirements can be verified prior to analysis using built-in preflight check functions in *ZonationR* (see below). For the workflows described here, we use layers provided by the *Zonation5RData* package, and manage the environment with the *withr* package (http://withr.r-lib.org/).

~~~
# Install the Zonation5RData package from GitHub:
pak::pak(“thiago-cav/Zonation5RData”)
# Load necessary libraries
library(ZonationR)
library(Zonation5RData)
# Create folder for biodiversity input data and copy example rasters
dir.create(“biodiversity”, showWarnings = FALSE)
files <-zonation5rdata_list(“biodiversity”, full.names = TRUE)
file.copy(files, “biodiversity”, overwrite = TRUE)
# Check Zonation executable
check_zonation_executable()
# Verify raster uniformity
check_raster_uniformity(“biodiversity”)
# Ensure working directory is writable
check_dir_writable(“.”)
~~~

Running the workflow inside an RStudio project is strongly recommended. It ensures consistent file paths, proper directory write permissions, and smooth execution of Zonation analyses. On Linux or WSL systems, some R package dependencies of *ZonationR*, particularly *terra*, require system development libraries (e.g., libgdal-dev, libgeos-dev, libproj-dev, libudunits2-dev) to compile successfully.

### 3.2 Single-variant prioritisation

Running a single prioritisation is straightforward. Users only need a folder with harmonized feature layers and the path to the Zonation installation directory. Once these are ready, a complete prioritisation can be run entirely within R with only a few lines of code:

~~~
# Create input files and run prioritisation
feature_list(spp_file_dir = “biodiversity”)
settings_file(feature_list_file = “feature_list.txt”)
command_file(zonation_path = “C:/Program Files (x86)/Zonation5”)
run_command_file(“.”)
~~~

This minimal example makes the workflow well suited for introductory teaching, as it allows users to focus on core concepts of spatial prioritisation without requiring complex setup or manual file handling. Outputs, including priority maps, feature curves, and performance curves, are saved in the output folder. This demonstrates that the first prioritisation variant can be obtained with minimal setup.

### 3.2 Multiple-variant prioritisation

*ZonationR* supports incremental variant workflows. Following Zonation best practices, analyses are developed incrementally through variants. Each variant modifies specific inputs or settings and is run in its own folder. This staged workflow keeps each variant isolated, reproducible, and easy to compare. In R, we use the package *withr* to temporarily change the working directory for each variant, keeping the project organized while running multiple scenarios. Folder names are based on key settings used in each scenario. For example, 03_cazmax_w indicates the third variant used the CAZMAX marginal loss rule with feature weights (Figure 2).

**Figure 2.**
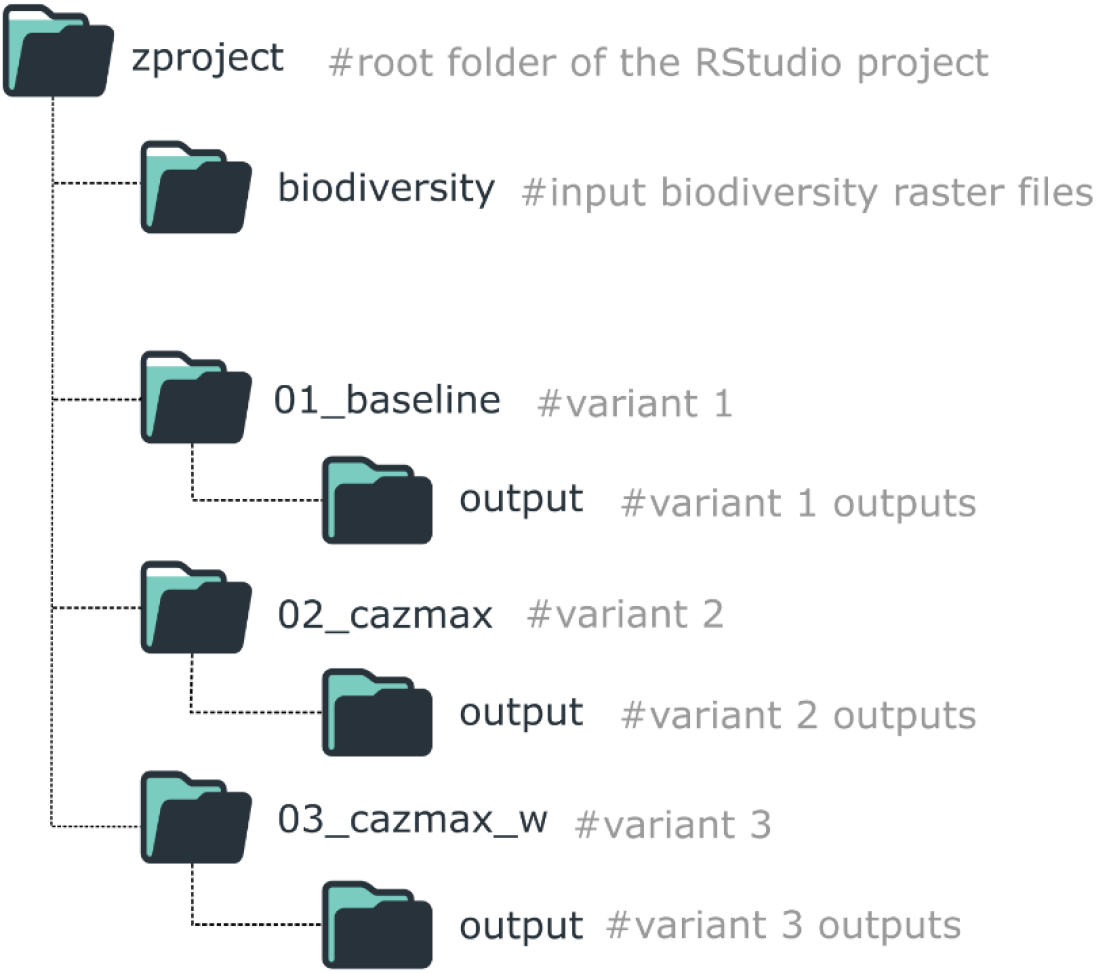
Folder structure for organizing a *ZonationR* workflow with multiple variants.

**Figure 3.**
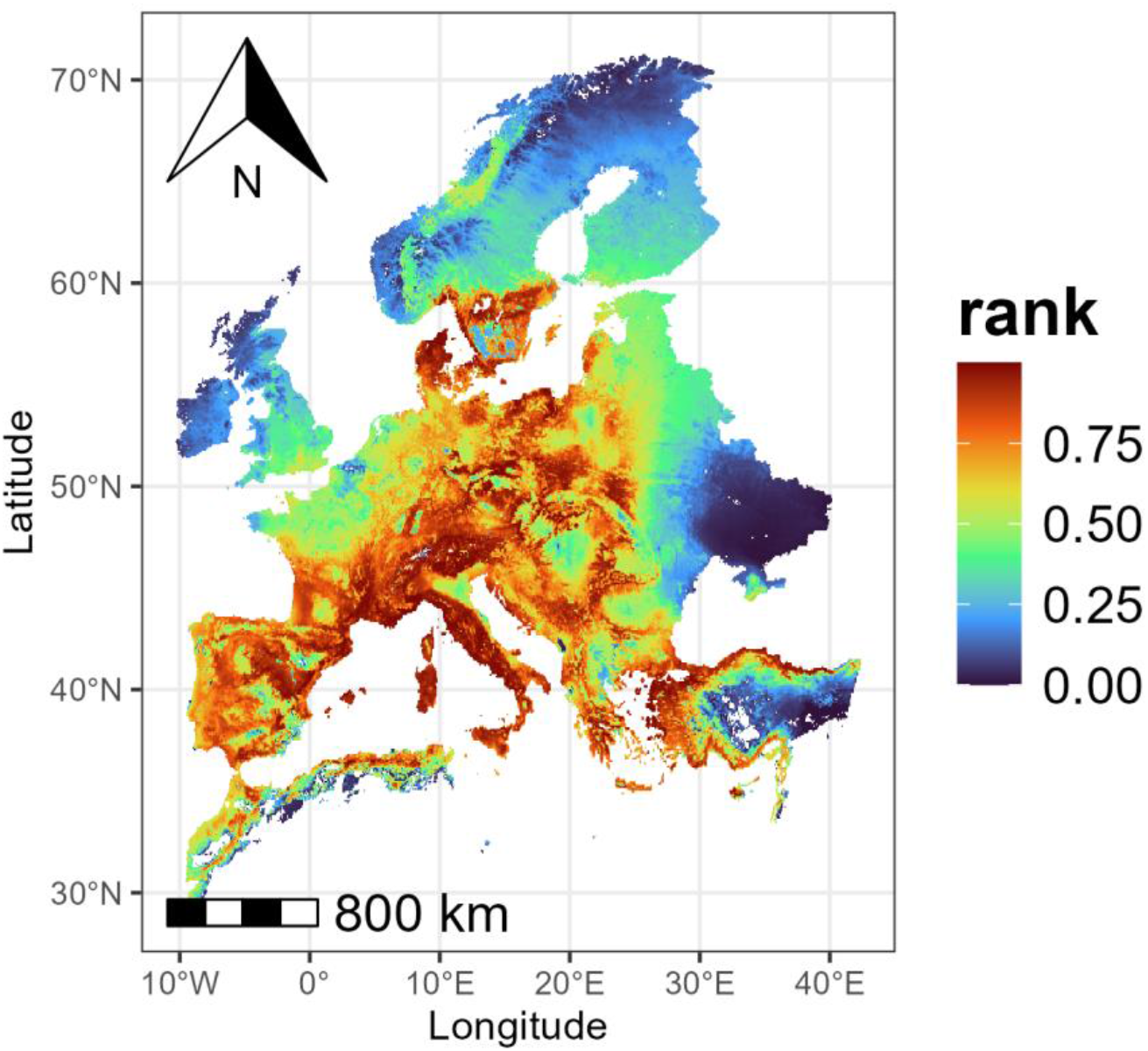
Example of a priority map generated with *ZonationR*. The function requires only the path to a workflow variant folder and produces a spatial map of conservation priorities, which can be customized using *ggplot2* syntax.

**Figure 4.**
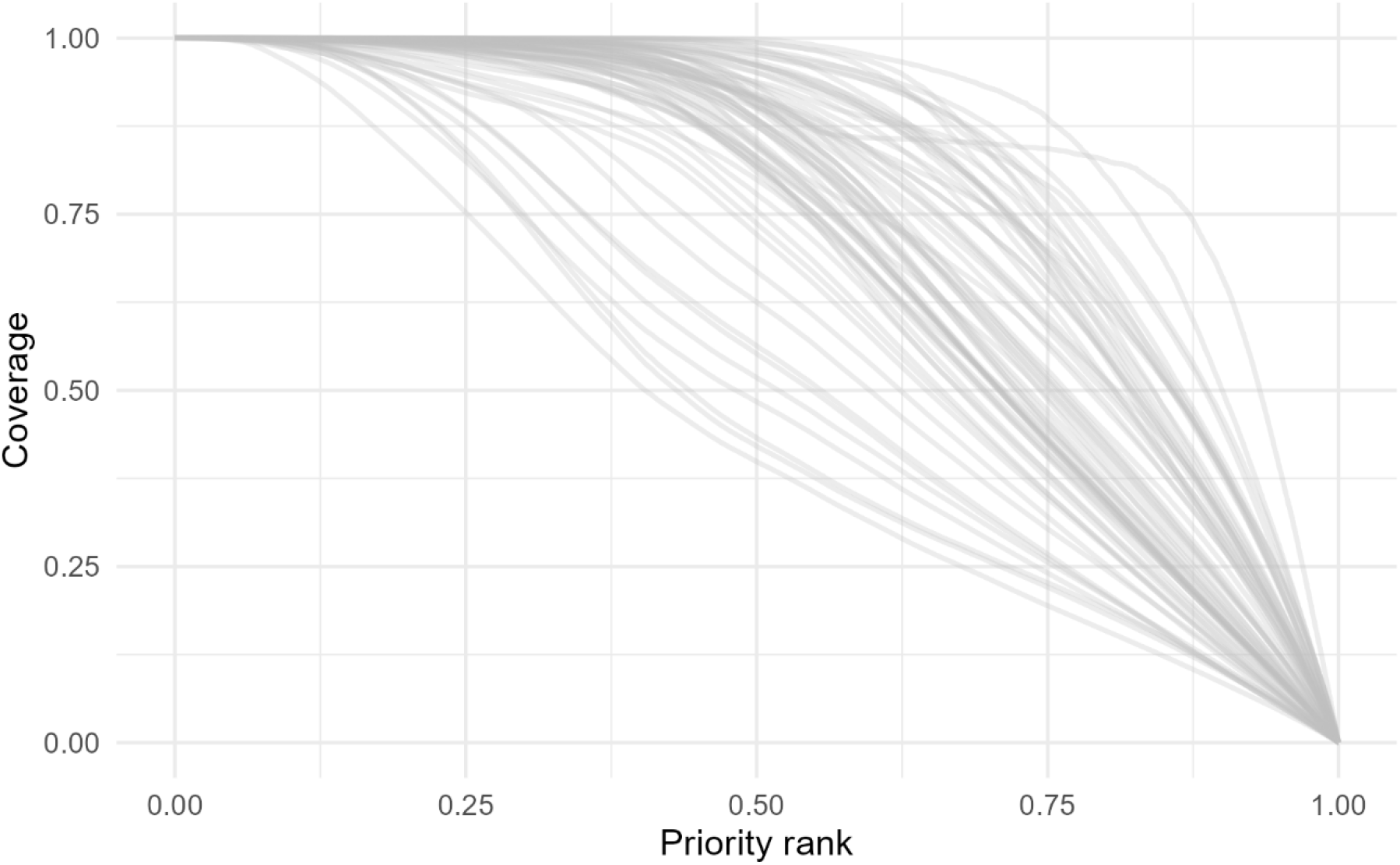
Feature-level performance curves generated with *ZonationR* from a workflow variant folder. These curves show how each feature is represented across the priority ranking.

#### Variant 1: Baseline

~~~
# Install ‘withr’ package if not already installed
if (!requireNamespace(“withr”, quietly = TRUE)) {
  install.packages(“withr”)
}
# Load necessary libraries
library(withr)
dir.create(“01_baseline”, showWarnings = FALSE)
withr::with_dir(“01_baseline”, {
  feature_list(spp_file_dir = “../biodiversity”)
  settings_file(feature_list_file = “feature_list.txt”)
  command_file(zonation_path = “C:/Program Files (x86)/Zonation5”)
  run_command_file(“.”)
})
~~~

#### Variant 2: CAZMAX marginal loss rule

In this variant, we change the marginal loss rule to Core-Area Zonation (CAZMAX). CAZMAX aims to always cover high-occurrence locations for all features, even if this comes at the cost of lower average coverage.

~~~
dir.create(“02_cazmax”, showWarnings = FALSE)
withr::with_dir(“02_cazmax”, {
  feature_list(spp_file_dir = “../biodiversity”)
  settings_file(feature_list_file = “feature_list.txt”)
  command_file(
    zonation_path = “C:/Program Files (x86)/Zonation5”,
    marginal_loss_mode = “CAZMAX”
)
  run_command_file(“.”)
})
~~~

#### Variant 3: CAZMAX with feature weights

We can assign unique weights to features (e.g., species) to increase the priority of areas where those features occur. This allows emphasizing key species or biodiversity features in the prioritisation. For this purpose, we can use the weight argument in the `feature_list()` function.

~~~
dir.create(“03_cazmax_w”, showWarnings = FALSE)
# setting weights
spp_files <-list.files(“biodiversity”, pattern = “\\.tif$“, full.names =
TRUE)
weights <-rep(1, length(spp_files))
weights[c(3,7,10)] <-5.0 # Prioritize selected species
withr::with_dir(“03_cazmax_w”, {
  feature_list(spp_file_dir = “../biodiversity”, weight = weights)
  settings_file(feature_list_file = “feature_list.txt”)
  command_file(
    zonation_path = “C:/Program Files (x86)/Zonation5”,
    marginal_loss_mode = “CAZMAX”,
    flags = “w”
)
  run_command_file(“.”)
})
~~~

This workflow demonstrates how to construct a variant-based prioritisation analysis using *ZonationR*, with each variant introducing new components to the prioritisation setup. Developing analyses in incremental variants improves transparency, reproducibility, and scenario comparison in spatial conservation planning.

### 3.3 Post-processing

*ZonationR* provides functions to explore and analyze outputs generated by Zonation. Users can visualize priority maps, inspect performance curves, and assess feature-level coverage, among other analytical functions (see Table 1). Its graphical functions leverage basic *ggplot2* syntax to produce customizable, publication-ready or exploratory figures, enabling straightforward evaluation of spatial prioritisation outcomes.

~~~
# Define the folder from the multiple variants workflow
variant_folder <-“01_baseline”
# Generate a priority map
priority <-priority_map(variant_folder)
print(priority)
# Generate feature-level performance curves
feature <-feature_curves(variant_folder)
print(feature)
~~~

Comprehensive guidance is provided on the package website (https://thiago-cav.github.io/ZonationR/), which includes all relevant documentation and articles (Table 5).

**Table 5.**
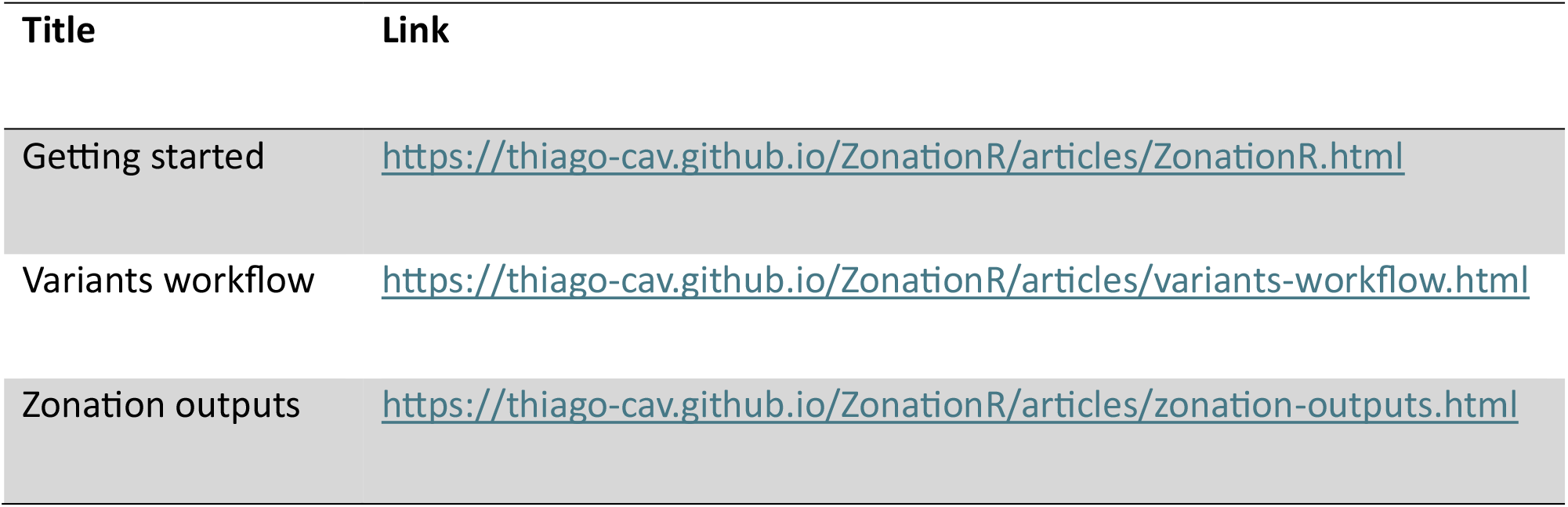
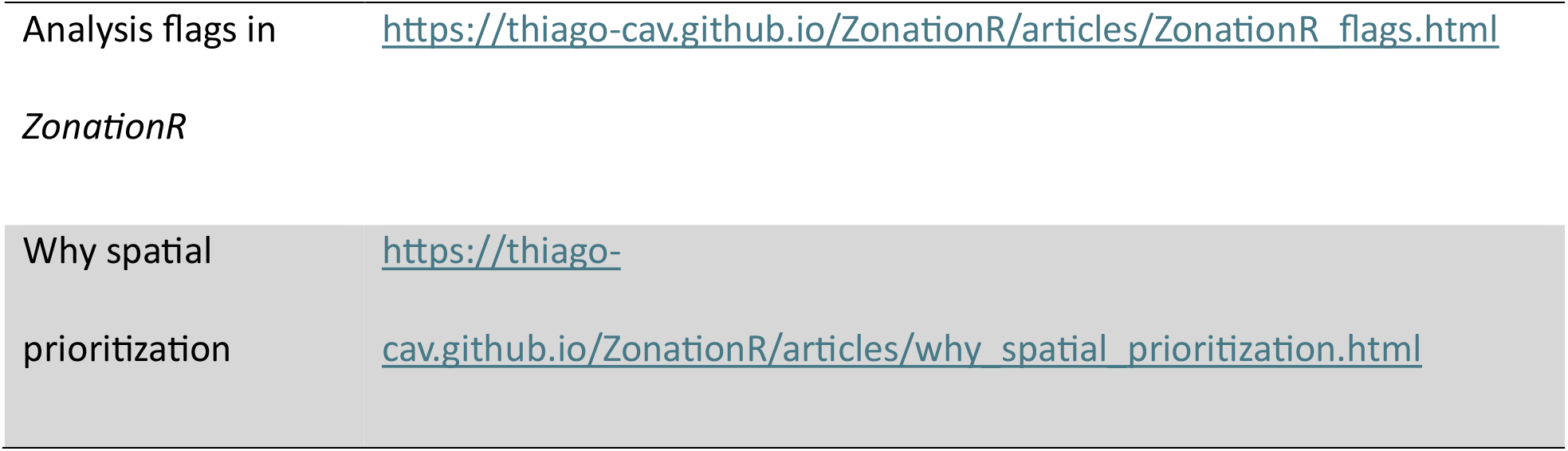
Overview of articles provided with the *ZonationR* package.

## 4. Conclusion

*ZonationR* provides an accessible R interface to the Zonation software, enabling users to perform spatial conservation prioritisation efficiently and reproducibly. The package supports both single- and multiple-variant analyses, allowing researchers to run prioritisations efficiently and compare scenarios. It also leverages built-in post-processing functions for visualisation, including priority maps, performance curves, and feature-level coverage plots. Beyond standard outputs such as rank maps and performance curves, *ZonationR* extends the workflow with additional post-processing tools. These enable, for instance, similarity assessments between prioritisation solutions, as well as analyses of cost accumulation and feature representation across landscapes. While not all functionalities of the Zonation software are currently implemented in *ZonationR*, the modular design makes it straightforward to edit the files created by the package to add any configuration or input files needed for other types of analyses. The package is actively maintained and will continue to be updated, with new functions and arguments added over time to accommodate additional Zonation parameters and use cases.

To support learning and teaching, the package includes comprehensive, user-friendly documentation with step-by-step tutorials, guiding users through setup, workflows, and interpretation of results, and making it suitable for both self-guided learning and formal training. Beyond these advantages, integrating Zonation with the R environment creates new opportunities for iterative exploration of conservation scenarios and systematic evaluation of methodological choices, as well as closer integration with broader ecological workflows (e.g., with species distribution modelling). As conservation planning increasingly relies on large, complex, multi-source datasets and integrative approaches, such integration is essential for enabling robust, transparent, and reproducible decision-making across spatial scales.

## Supporting information

Articles demonstrating the application of ZonationR

Articles demonstrating the application of ZonationR

Additional information on the data used as an example in this paper

## Acknowledgements

This research was supported by the NaturaConnect project, funded by the European Union’s Horizon Europe research and innovation programme (grant agreement No. 101060429).

## Author Contributions

Thiago Cavalcante conceptualised the study and led the design of the software package. The software was developed with input from Bruno Ribeiro, Karlo Guidoni-Martins, and Heini Kujala. Thiago Cavalcante led the writing of the manuscript and wrote the R package documentation.

All authors contributed critically to the drafts and gave final approval for publication.

## Data availability statement

*ZonationR* source code is available on GitHub at https://github.com/thiago-cav/ZonationR. A companion data package, *Zonation5RData*, is also available on GitHub at https://github.com/thiago-cav/Zonation5RData and provides all example GeoTIFF layers required to reproduce the workflows demonstrated in this manuscript. Details on the original data sources used to construct these example datasets are provided in the package documentation (README) of *Zonation5RData*. All analyses are based on these example datasets and publicly available software.

## References

Arponen, A., Heikkinen, R. K., Thomas, C. D. and Moilanen, A. 2005. The value of biodiversity in reserve selection: representation, species weighting, and benefit functions. Conservation Biology, 19, 2009–2014.

Cavalcante, T., Kujala, H., Virtanen, E. A., O’Connor, L., Lehtinen, P. and Moilanen, A. 2025. Evaluating trade-offs between species targets and average coverage in spatial conservation planning. Biological Conservation, 310, 111368. 10.1016/j.biocon.2025.111368

Giakoumi, S. et al. 2025. Advances in systematic conservation planning to meet global biodiversity goals. Trends in Ecology and Evolution, 40, 395-410. 10.1016/j.tree.2024.12.002

Lehtomäki, J. and Moilanen, A. 2013. Methods and workflow for spatial conservation prioritization using Zonation. Environmental Modelling and Software, 47, 128–137.

Margules, C. R. and Pressey, R. L. 2000. Systematic conservation planning. Nature, 405, 243–253.

Moilanen, A., Franco, A. M., Early, R. I., Fox, R., Wintle, B. and Thomas, C. D. 2005. Prioritizing multiple-use landscapes for conservation: methods for large multi-species planning problems. Proceedings of the Royal Society B: Biological Sciences, 272, 1885–1891.

Moilanen, A., Wilson, K. and Possingham, H. 2009. Spatial conservation prioritization: quantitative methods and computational tools: Oxford University Press.

Moilanen, A., Lehtinen, P., Kohonen, I., Jalkanen, J., Virtanen, E. A. and Kujala, H. 2022. Novel methods for spatial prioritization with applications in conservation, land use planning and ecological impact avoidance. Methods in Ecology and Evolution, 13, 1062-1072. 10.1111/2041-210X.13819

Sarkar, S., Pressey, R. L., Faith, D. P., Margules, C. R., Fuller, T., Stoms, D. M., Moffett, A., Wilson, K. A., Williams, K. J. and Williams, P. H. 2006. Biodiversity conservation planning tools: present status and challenges for the future. Annual Review of Environment and Resources, 31, 123–159.

Wickham, H. and Bryan, J. 2023. R packages: O’Reilly Media, Inc.

