## Supplementary material for "ZonationR: An R interface to the Zonation software for reproducible spatial conservation prioritisation workflows": Articles demonstrating the application of ZonationR

### Working with variants in ZonationR

Source: vignettes/articles/variants-workflow.Rmd

This vignette demonstrates how to explore multiple prioritization scenarios using *ZonationR*. Users will:

1. Run a baseline prioritization.
2. Modify the marginal loss rule.
3. Apply feature weights.
4. Restrict the analysis to a geographic mask.
5. Include cost considerations.

Following Zonation best practices, analyses are developed incrementally through variants. Each variant modifies specific inputs or settings and is run in its own folder. Folder names are based on key settings used in each scenario. For example, **03_cazmax_w** indicates the third variant used the CAZMAX marginal loss rule with feature weights:

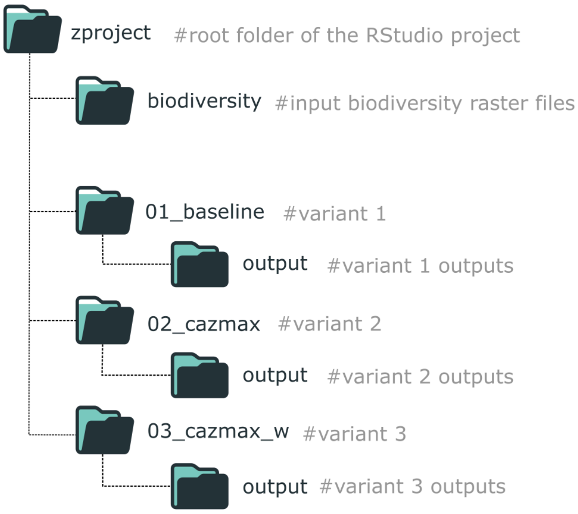


Folder structure for organizing a *ZonationR* workflow with multiple variants.

This staged workflow keeps each variant **isolated, reproducible, and easy to compare**. In R, we use `withr::with_dir()` to temporarily change the working directory for each variant, keeping the project organized while running multiple scenarios.

#### Setup

```
  install.packages("withr")
}

# Load necessary libraries
library(ZonationR)
library(Zonation5RData)
library(withr)
```

#### Prepare input data

In this tutorial, we use the Zonation5RData package, which provides GeoTIFF layers for workflows with Zonation and *ZonationR*. To use the layers in the **RStudio project**, we first create folders and then copy the raster files from the package.

```
# Create folders to store the input data
dir.create("biodiversity", showWarnings = FALSE)
dir.create("other_layers", showWarnings = FALSE)

# Get the paths to biodiversity rasters included in the package
files_biod <- zonation5rdata_list("biodiversity", full.names = TRU

# Copy the biodiversity rasters to the local biodiversity folder
file.copy(files_biod, "biodiversity", overwrite = TRUE)

# Get the paths to the additional layers included in the package
files_other <- zonation5rdata_list("other_layers", full.names = TI

# Copy these rasters to the local other_layers folder
file.copy(files_other, "other_layers", overwrite = TRUE)
```

#### Variant 1

We begin with a baseline prioritization using default settings.

```
# Run baseline variant in folder 01_baseline
dir.create("01_baseline", showWarnings = FALSE)
withr::with_dir("01_baseline", {
  feature_list(spp_file_dir = "../biodiversity")
  settings_file(feature_list_file = "feature_list.txt")
  command_file(zonation_path = "C:/Program Files (x86)/Zonation5"
  run_command_file(".")
})
```

(CAZMAX). CAZMAX aims to always cover high-occurrence locations for all features, even if this comes at the cost of lower average coverage.

```
# Create the folder
dir.create("02_cazmax", showWarnings = FALSE)

withr::with_dir("02_cazmax", {
  feature_list(spp_file_dir = "../biodiversity")
  settings_file(feature_list_file = "feature_list.txt")
  command_file(zonation_path      = "C:/Program Files (x86)/Zonat
               marginal_loss_mode = "CAZMAX" # We edit the margin
  )
  run_command_file(".")
})
```

##### Variant 3

We can assign unique weights to features (e.g., species) to increase the priority of areas where those features occur. This allows emphasizing key species or biodiversity features in the prioritization. For this purpose, we use the `weight` argument in the `feature_list()` function. To ensure that these weights are applied during analysis, the `w` flag must also be included in the `command_file()` call.

> For a complete reference of all available analysis flags and their requirements in ZonationR, please see the vignette Analysis flags in ZonationR.

```
# Create the folder
dir.create("03_cazmax_w", showWarnings = FALSE)

# List files from the folder
spp_files <- list.files("biodiversity", pattern = "\\.tif$", full

# Create a weight vector: default weight = 1 for all species
weights <- rep(1, length(spp_files))

# Set higher weights for selected key species
# For example, species 3, 7, and 10 are more important
weights[c(3, 7, 10)] <- 5.0

# Run the prioritization
withr::with_dir("03_cazmax_w", {
```

```
  command_file(zonation_path = "C:/Program Files (x86)/Zonation5"
               marginal_loss_mode = "CAZMAX",
               flags = "w" # Flag w (use weights)
  )
  run_command_file(".")
})
```

#### Variant 4

In this variant, we restrict the prioritization to a subset of European countries. Only areas within the area mask are considered for the ranking process. This approach is useful for focused conservation planning in a defined geographic region.

```
dir.create("04_cazmax_wa", showWarnings = FALSE)

withr::with_dir("04_cazmax_wa", {
  feature_list(spp_file_dir = "../biodiversity",
               weight = weights)
  settings_file(feature_list_file = "feature_list.txt",
                analysis_area_mask_layer = "../other_layers/area_
  command_file(zonation_path = "C:/Program Files (x86)/Zonation5"
               marginal_loss_mode = "CAZMAX",
               flags = "wa" # Flag a for analysis area mask
  )
  run_command_file(".")
})
```

#### Variant 5

In this variant, we include a cost layer using the Global Human Modification (GHM) dataset. Cells with higher human impact are penalized, reducing their priority in the ranking.

```
dir.create("05_cazmax_waX", showWarnings = FALSE)

withr::with_dir("05_cazmax_waX", {
  feature_list(spp_file_dir = "../biodiversity",
               weight = weights)
  settings_file(feature_list_file = "feature_list.txt",
                analysis_area_mask_layer = "../other_layers/area_
                cost_layer = "../other_layers/gHM_Europe.tif") # 
  command_file(zonation_path = "C:/Program Files (x86)/Zonation5"
```

```
  run_command_file(".")
})
```

#### Key takeaways

This workflow demonstrated how to construct a **variant-based prioritization analysis** using *ZonationR*, with each variant introducing new components to the prioritization setup:

| Variant | Description |
|---|---|
| | settings |
| 02 | Prioritization with CAZMAX marginal loss rule |
| 03 | CAZMAX rule plus feature/species weighting applied |
| 04 | CAZMAX with weighting and restricted to a geographic mask |
| 05 | CAZMAX with weighting, geographic mask, and cost layer (GHM) |

Developing analyses in incremental variants improves **transparency, reproducibility, and scenario comparison** in spatial conservation planning. To explore the outputs generated by these analyses (e.g., rank maps, performance curves), see the Zonation outputs vignette, which provides guidance on interpreting and visualizing Zonation results.

#### On this page

Setup

Prepare input data

Variant 1

Variant 2

Variant 3

Key takeaways

Developed by Thiago Cavalcante, Bruno Ribeiro, Karlo Guidoni-Martins, Heini Kujala.

Site built with pkgdown 2.2.0.
