## Supplementary material for "ZonationR: An R interface to the Zonation software for reproducible spatial conservation prioritisation workflows": Articles demonstrating the application of ZonationR

### Exploring Zonation outputs

Source: vignettes/articles/zonation-outputs.Rmd

This vignette demonstrates how to explore and interpret some of the main outputs generated by Zonation using the *ZonationR* package. Specifically, users will:

1. Examine spatial patterns of conservation priorities.
2. Inspect performance curves.
3. Assess the distribution of feature coverage.

#### Setup

##### Installation

```
# Install required packages if not already installed
if (!requireNamespace("ggplot2", quietly = TRUE)) {
  install.packages("ggplot2")
}

if (!requireNamespace("patchwork", quietly = TRUE)) {
  install.packages("patchwork")
}
```

##### Libraries

```
# Load necessary libraries
library(ZonationR)
library(Zonation5RData)
library(ggplot2)
library(patchwork)
```

#### Prepare input data

Here, we use the baseline variant output folder from the vignette Variants worflow to illustrate some of the post-processing functions. You can either use the output folder data provided by the Zonation5RData package, as shown below, or, if you are following the multiple variants tutorial, you can use any output folder from there.

```
# Copy the whole directory, preserving the structure
file.copy(src, ".", recursive = TRUE, overwrite = TRUE)

# Define a variable for the baseline folder
baseline_folder <- "01_baseline"
```

#### Priority map

One of the main outputs of Zonation is the priority rank map, which shows the conservation priority of each cell across the landscape. Higher numbers indicate areas that are more important for conservation, while lower numbers indicate areas of lower priority. Each rank value tells you how a cell compares to others in terms of conservation priority. But please note that **the scale is relative, not absolute**. For example, a cell ranked 0.8 is not twice as valuable as one ranked 0.4. The priority map should then be considered together with other outputs, like performance curves, to fully understand the spatial distribution of conservation value, potential trade-offs, and the conservation effectiveness. We can use the `priority_map()` function to visualize the ranking values across the landscape:

```
p1 <- priority_map(baseline_folder)

print(p1)
```

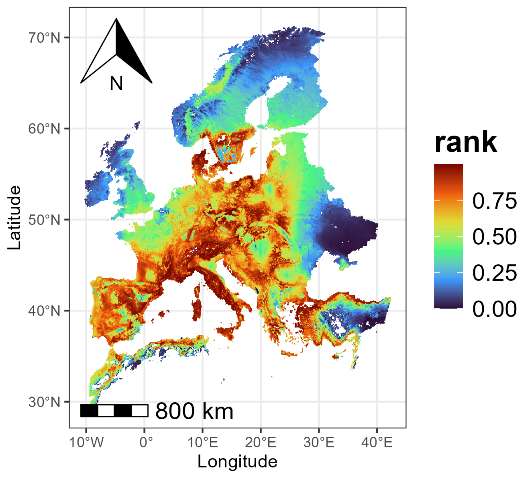


In addition to visualizing the continuous priority values, we can also adjust the map to make important areas stand out more clearly. The `classify` argument in `priority_map()` allows choosing whether to display the map as continuous or divided into classes. If `classify = TRUE`, the map is divided into discrete classes, and we can define custom breaks

```
breaks <- c(0, 0.1, 0.5, 0.9, 1)
labels <- c("very low", "low", "medium", "very high")
p2 <- priority_map(baseline_folder, classify = TRUE,
                   breaks = breaks, labels = labels)
print(p2)
```

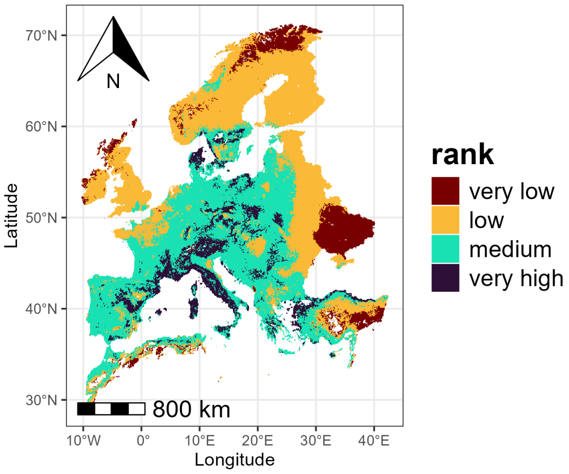


The visualization functions in *ZonationR* return `ggplot2` objects, which means that users can further customize the plots using the `ggplot2` ecosystem. For example, we can apply a different color palette to the map:

```
p2 <- p2 + scale_fill_viridis_d() + labs(fill = "rank")
print(p2)
```

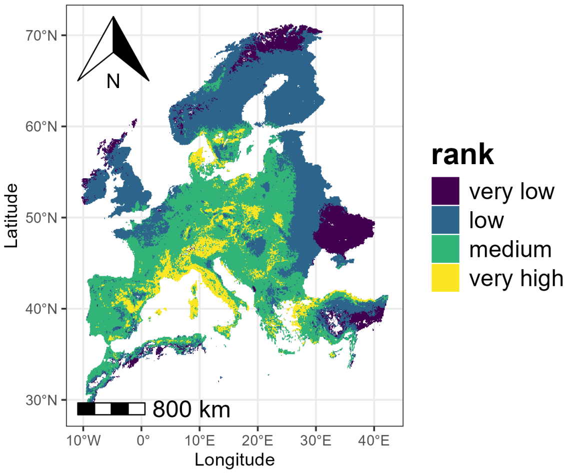


#### Performance curves

The performance curves describe how much of each feature's distribution would be covered if we protected grid cells following their priority rank order. The `summary_curves()` function is helpful for understanding

- **mean** - average performance across all species
- **max** - performance of the best-performing species
- **min** - performance of the worst-performing species

```
p3 <- summary_curves(baseline_folder, metrics = c("mean", "max", 
  ggtitle("Summary Curves") +
  theme(plot.title = element_text(hjust = 0.5))

print(p3)
```

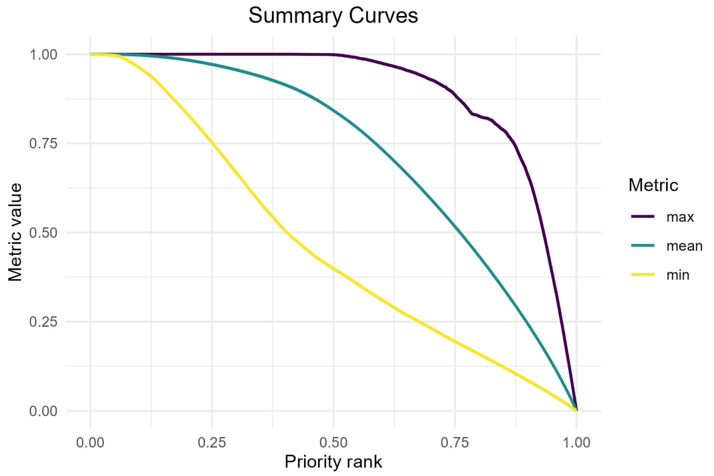


We can also visualize additional metrics from the summary curves file using separate panels (facets). The `facet` argument should be set to `TRUE` when plotting metrics that have different units or value ranges.

```
p4 <- summary_curves(baseline_folder,
                     metrics = c("remaining_area", "mean"),
                     facet = TRUE)
print(p4)
```

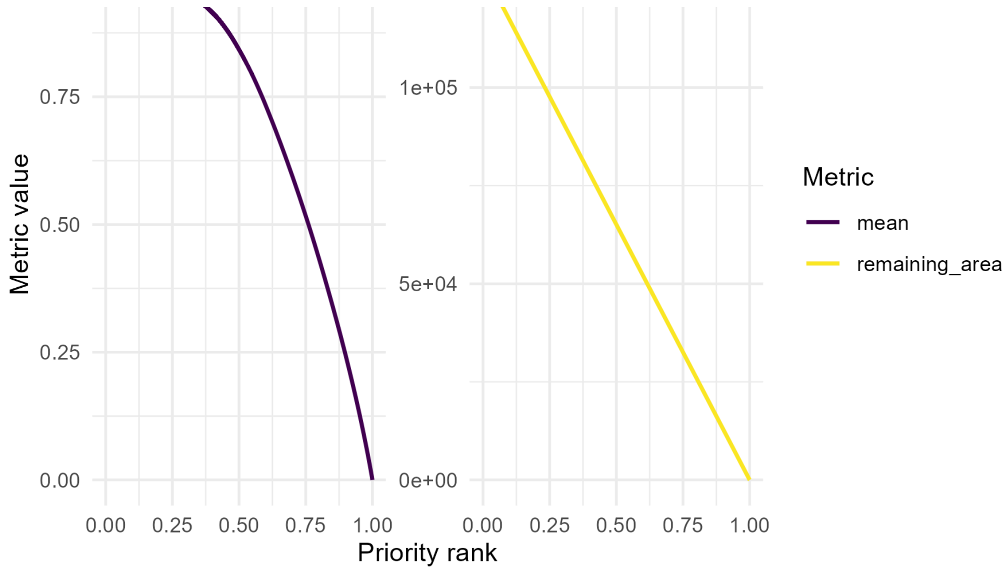


Besides looking at overall performance with `summary_curves()`, one could also be interested in how each feature is represented across the priority ranking, which can be explored using the `feature_curves()` function:

```
p5 <- feature_curves(baseline_folder) +
  ggtitle("Feature curves") +
  theme(plot.title = element_text(hjust = 0.5))

print(p5)
```

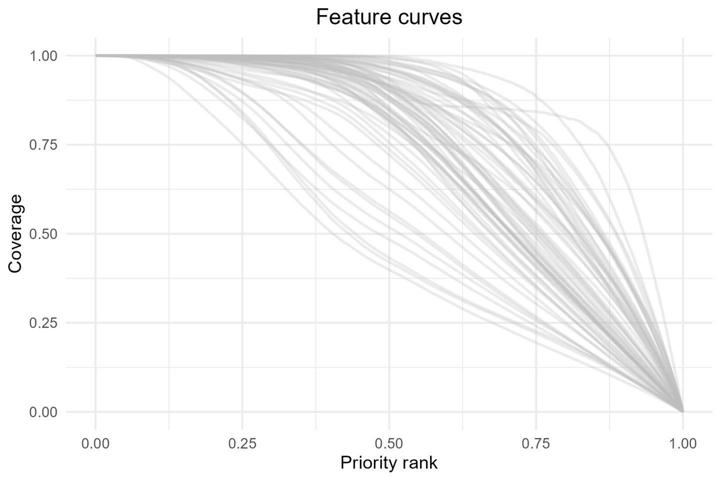


As with `priority_map()`, the functions `summary_curves()` and `feature_curves()` return `ggplot2` objects, allowing further customization of the plots. For example, we can add a vertical line to highlight the top 10% ( `rank = 0.9` ) priority cells.

```
p5 <- p5 + geom_vline(xintercept = 0.9, linetype = "dashed", colo
```

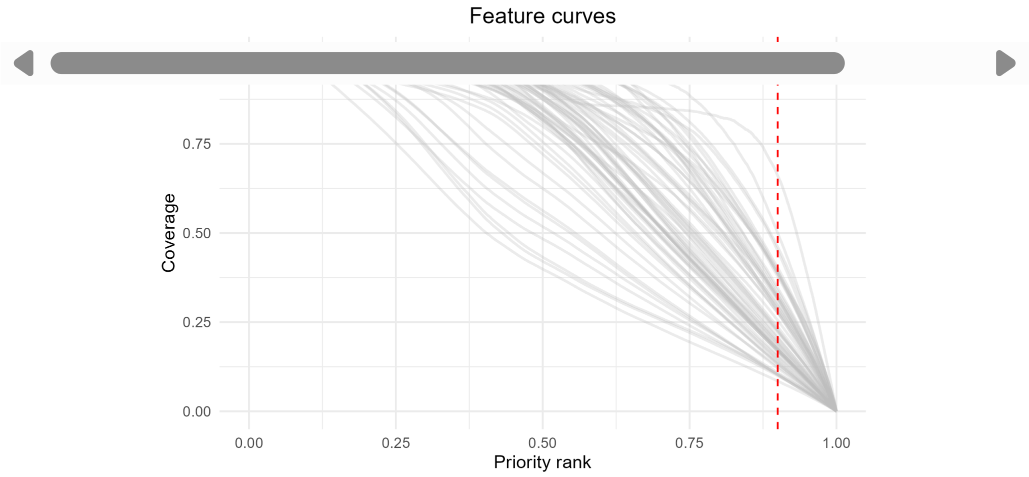


> The vertical line at 0.9 represents the top 10% priority areas. Intersection with the curves shows the proportion of each feature representation within these areas.

As mentioned earlier, priority rank maps and performance curves should be interpreted together. For example, we could use the `priority_map()` function to plot a classified map showing only the top 10% priority areas, alongside the corresponding feature curves.

```
# Define the threshold for the top 10% priority cells
threshold <- 0.90

# Define breaks and labels for classified map
breaks <- c(0, 0.90, 1)              # 0-90% = other areas, 90-100%
labels <- c("other areas", "top 10%")

# Plot the classified map with two colors
top10_map <- priority_map(
  baseline_folder,
  classify = TRUE,
  breaks = breaks,
  labels = labels
) +
  scale_fill_manual(values = c("lightgray", "darkgreen")) +
  ggtitle("Priority map") +
  theme(plot.title = element_text(hjust = 0.5),
        legend.position = "bottom") +
  labs(fill = "rank")

# Combine map and feature curves
p1_combined <- top10_map + p5
```

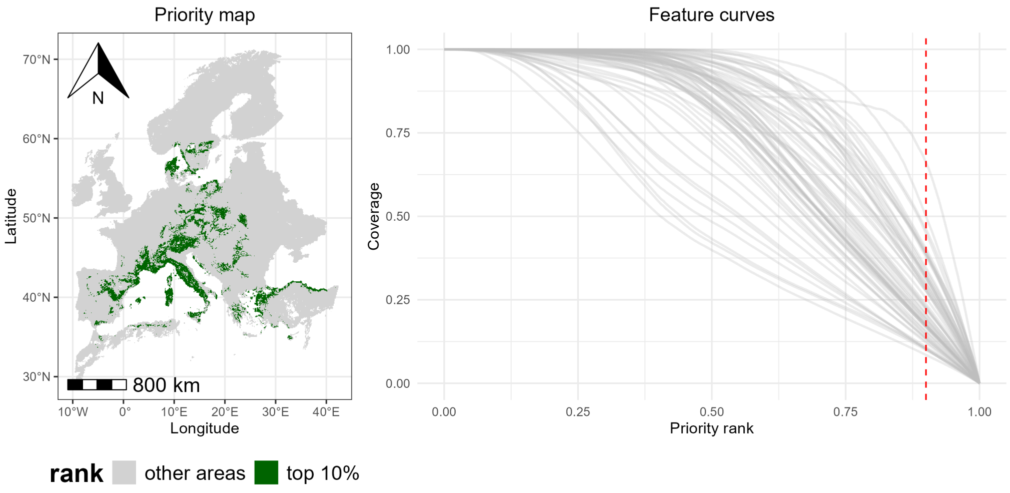


By combining the classified map and the feature curves side by side, we can see which areas are top-priority and how well each species is represented within those areas (i.e., the curves intersecting the red dashed line), providing a integrated view of spatial priorities and species coverage.

#### Coverage distribution

In addition to maps and performance curves, we can also explore the coverage distribution for any selected priority fraction using the function `coverage_distribution()`. The histogram shows, for each coverage level (or coverage bin) on the x-axis, the number of species reaching that coverage level (y-axis). For example, a bar reaching 30 on the y-axis at the coverage range 0.1-0.2 means that 30 species have between 10% and 20% of their range included in the selected priority fraction.

```
p6 <- coverage_distribution(baseline_folder, target_rank = 0.9) +
  ggtitle("Coverage at the top 10%") +
  theme(plot.title = element_text(hjust = 0.5))

print(p6)
```

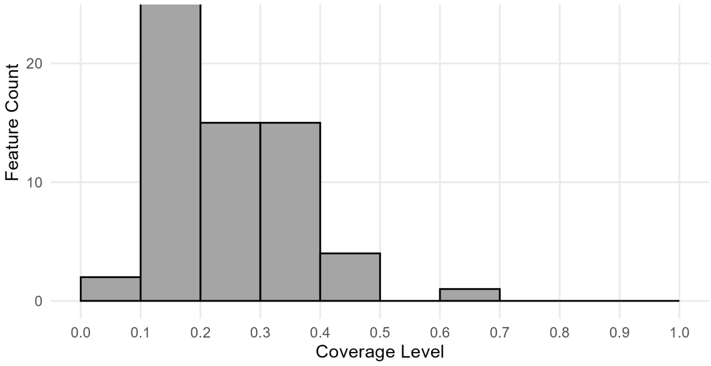


> An overview of Zonation results can be created by combining maps and other outputs.

```
p2_combined <- top10_map + p5 / p6

print(p2_combined)
```

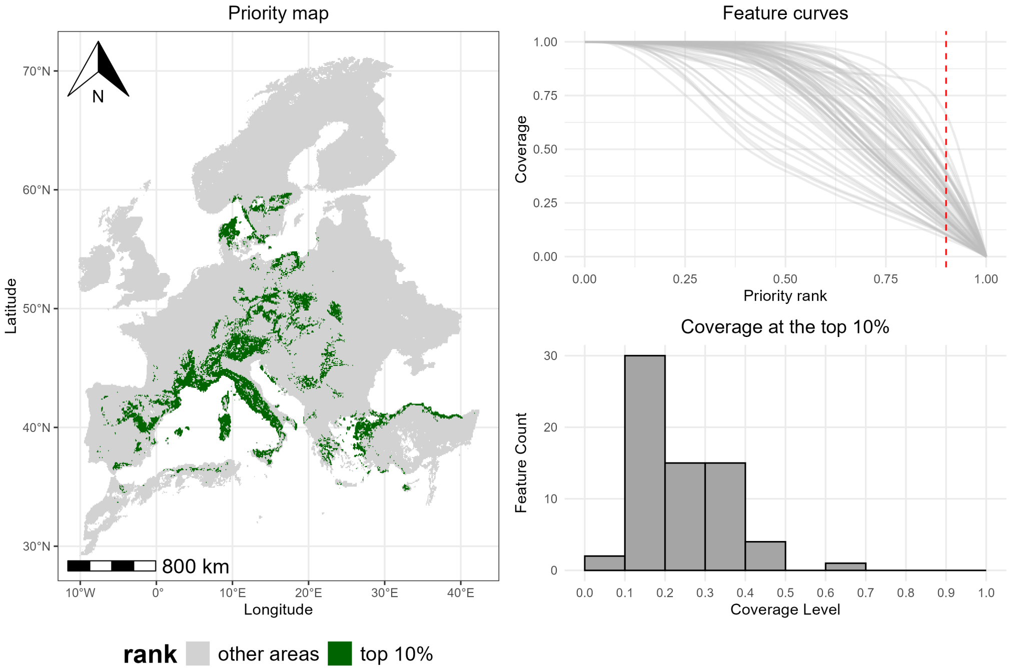


#### Key takeaways

By looking at the combined visualization, we can see which areas are top-priority (**map**), how well each species is represented in those areas (**feature curves**), and the number of species at different coverage levels (**histogram**). This provides a full view of spatial priorities and feature-level coverage, illustrating how *ZonationR* allows users to explore and interpret conservation outcomes in a flexible and straightforward way.

ZonationR 1.0.1.9000

- Setup
  - Installation
  - Libraries
- Prepare input data
- Priority map
- Performance curves
- Coverage distribution
- Key takeaways

---

Developed by Thiago Cavalcante, Bruno Ribeiro, Karlo Guidoni-Martins, Heini Kujala.

Site built with pkgdown 2.2.0.
