## Additional information on the data used as an example in this paper for "ZonationR: An R interface to the Zonation software for reproducible spatial conservation prioritisation workflows"

### Zonation5RData Overview

#### Directory layout (in `inst/extdata/`)

| Path | Contents |
|---|---|
| `biodiversity/` | EU-Trees4F SDM rasters (one GeoTIFF per species; filenames use underscore-separated Latin names). |
| `other_layers/` | Cost ( `gHM_Europe.tif` , gHM-derived) and planning mask ( `area_mask.tif` ). |
| `01_baseline/` | Output files from a baseline Zonation prioritization analysis. |
| `metadata.md` | This file: sources and attribution. |

This package contains a curated dataset designed for educational purposes within the Zonation software framework. The dataset has been processed and organized to facilitate learning and experimentation, using freely available data that are fully referenced. The main objective is to provide an accessible resource for educators and students to explore *ZonationR* capabilities and deepen their understanding of spatial conservation planning. Please note that this dataset (or parts of it) may not fully align with real-world research requirements. Please acknowledge this package and all included materials when using them in your work. All raster layers are stored as GeoTIFF files and are georeferenced to the WGS84 coordinate reference system (EPSG:4326).

### 1. Biodiversity Features

The biodiversity folder contains species distribution models (SDMs) retrieved from the **EU-Trees4F** dataset (Mauri et al., 2022). This dataset provides SDMs for 67 tree species across Europe at a 10 km spatial resolution. The predictions were constrained to account for ecological factors such as dispersal limitations.

### 2. Other Layers

This folder contains supporting spatial data used in Zonation analyses, such as cost or mask layers, which provide information on constraints, planning costs, or the spatial extent to guide and refine the prioritization of conservation areas.

#### 2.1 Cost Layer

The gHM layer is derived from the **Global Human Modification (gHM)** (https://developers.google.com/earth-engine/datasets/catalog/CSP_HM_GlobalHumanModification) dataset, which provides a comprehensive measure of human alteration of terrestrial lands worldwide, originally at 1 km² resolution. Values range from 0 to 1.0, representing the proportion and intensity of human modification for specific stressors (e.g., human settlement, agriculture, infrastructure). The data were retrieved and processed using Google Earth Engine.

#### 2.2 Area Mask

The area mask layer was created by rasterizing the boundaries of European countries, excluding certain overseas territories (e.g., the Canary Islands and the Azores). This layer defines the spatial extent for the study, ensuring analyses are limited to the target regions.

### 3. Baseline Zonation Output

The `01_baseline` folder contains output files generated from a baseline Zonation prioritization analysis using the biodiversity and supporting layers included in this package. The folder includes files such as rank maps, performance curves, and summary statistics produced by the Zonation software.

### References

Mauri, Achille, Marco Girardello, Giovanni Strona, Pieter SA Beck, Giovanni Forzieri, Giovanni Caudullo, Federica Manca, and Alessandro Cescatti. 2022. “EU-Trees4F, a Dataset on the Future Distribution of European Tree Species.” Journal Article. *Scientific Data* 9 (1): 37.
